# Host-induced gene silencing compromises Verticillium wilt in tomato and *Arabidopsis*

**DOI:** 10.1101/076976

**Authors:** Yin Song, Bart P.H.J. Thomma

## Abstract

Verticillium wilt, caused by soil-borne fungi of the genus *Verticillium*, is an economically important disease that affects a wide range of host plants. Unfortunately, host resistance against Verticillium wilts is not available for many plant species, and the disease is notoriously difficult to combat. Host-induced gene silencing (HIGS) is an RNA interference (RNAi) based process in which small RNAs are produced by the host plant to target parasite transcripts. HIGS has emerged as a promising strategy for improving plant resistance against pathogens by silencing genes that are essential for these pathogens. Here, we assessed whether HIGS can be utilized to suppress Verticillium wilt disease by silencing previously identified virulence genes of *V. dahliae* through the host plants tomato and *Arabidopsis*. In transient assays, tomato plants were agroinfiltrated with *Tobacco rattle virus* (TRV) constructs to target *V. dahliae* transcripts. Subsequent *V. dahliae* inoculation revealed suppression of Verticillium wilt disease in some, but not all, cases. Next, expression of RNAi constructs targeting *V. dahliae* transcripts was pursued in stable transgenic *Arabidopsis thaliana* plants. Also in this host, *V. dahliae* inoculation revealed reduced Verticillium wilt disease in some cases. Thus, our study suggests that, depending on the target gene chosen, HIGS against *V. dahliae* is operational in tomato and *A. thaliana* plants and may act as a plant protection approach that may be used in Verticillium wilt-susceptible crops.

## INTRODUCTION

Verticillium wilts are vascular wilt diseases that are caused by soil-borne fungi of the genus *Verticillium* (Fradin and Thomma, 2006; Klimes et al., 2015). This genus comprises ten species of soil-borne fungi that differ in their morphological features, such as resting structures, as well as in their ability to cause plant diseases (Inderbitzin et al., 2011). Within the *Verticillium* genus, *V. dahliae* is the most notorious pathogenic species that can infect hundreds of dicotyledonous hosts, including ecologically important plants and many high-value crops worldwide (Fradin and Thomma, 2006; Klosterman et al., 2009; Inderbitzin et al., 2011). Verticillium wilt diseases are difficult to control due to the long viability of the resting structures, the wide host range of the pathogens, and the inability of fungicides to affect the pathogen once in the plant vascular system. Thus, the most sustainable way to control Verticillium wilt diseases is the use of resistant cultivars. Polygenic resistance to *Verticillium* spp. has been described for several plant species, including potato, hop, alfalfa, cotton and strawberry (Simko et al., 2004; Bolek et al., 2005; Wang et al., 2008; Yang et al., 2008; Jakse et al., 2013; Antanaviciute et al., 2015), whereas single dominant resistance genes have been identified only in tomato, sunflower, cotton, potato and lettuce (Schaible et al., 1951; Putt, 1964; Barrow, 1970; Lynch et al., 1997; Mert et al., 2005; Hayes et al., 2011; Christopoulou et al., 2015). In tomato (*Solanum lycopersicum*), a single dominant locus that confers *Verticillium* resistance has been identified as the *Ve* locus, which controls *Verticillium* isolates that are assigned to race 1, whereas race 2 strains escape recognition (Schaible et al., 1951; Pegg, 1974). The *Ve* locus contains two closely linked and inversely oriented genes, *Ve1* and *Ve2*, both of which encode extracellular leucine rich repeat (eLRR) receptor-like proteins (RLPs) (Kawchuk et al., 2001; Wang et al., 2010). Of these, only *Ve1* was found to confer resistance against race 1 isolates of *Verticillium* in tomato (Fradin et al., 2009). Interestingly, interfamily transfer of *Ve1* from tomato to *Arabidopsis thaliana* has resulted in race-specific *Verticillium* resistance in the latter species (Fradin et al., 2011; 2014; Zhang et al., 2014), implying that the underlying immune signaling pathway is conserved (Fradin et al., 2011; Thomma et al., 2011). Tomato Ve1 serves as an immune receptor for recognition of the effector protein Ave1 that is secreted by race 1 strains of *V. dahliae* (de Jonge et al., 2012). More recently, homologs of tomato Ve1 acting as immune receptors that govern resistance against *V. dahliae* race 1 strains through recognition of the Ave1 effector have been characterized in other plant species including tobacco, potato, wild eggplant and hop, suggesting an ancient origin of the immune receptor Ve1 (Song et al., 2016).

Although the tomato *Ve1* gene is still currently deployed in tomato cultivars, isolates of *Verticillium* that escape Ve1-mediated recognition appeared within a few years after the introduction of the tomato *Ve1* (Pegg and Brady, 2002). These race 2 isolates of *Verticillium* steadily supplanted race 1 strains in various regions because of the extensive use of *Verticillium* race 1-resistant cultivars (Dobinson et al., 1996). So far, there remains no source of commercially employed resistance to *Verticillium* race 2 strains.

RNA interference (RNAi) is a conserved regulatory mechanism that affects gene expression in eukaryotic organisms (Baulcombe, 2005). RNA silencing is triggered by the processing of double stranded RNA (dsRNA) precursors into short interfering RNA (siRNAs) duplexes of 21-28 nucleotides in length, and followed by the guided cleavage or translational repression of sequence-complementary single-stranded RNAs by the generated siRNAs duplexes, which are incorporated into a silencing complex called RISC (RNA-induced silencing complex) (Ruiz-Ferrer and Voinnet, 2009). Plants and other eukaryotes have evolved RNAi machineries that not only regulate developmental programs, but also provide protection from invaders, such as viruses. In plants, RNAi has been exploited extensively and has become a powerful functional genomics tool to silence the expression of genes of interest as well as to engineer viral resistance (Duan et al., 2012). Interestingly, organisms that live within, or develop intimate contact with, a host, such as bacteria (Escobar et al., 2001; Escobar et al., 2002), nematodes (Huang et al., 2006), insects (Baum et al., 2007; Mao et al., 2007) and parasitic plants (Tomilov et al., 2008), are sensitive to small RNAs generated by the host and that are targeted to parasite transcripts. This so-called host-induced gene silencing (HIGS) has also emerged as a promising strategy against plant pathogens, including fungi and oomycetes. Initial reports of HIGS against filamentous pathogens were described for the maize kernel and ear rot pathogen *Fusarium verticillioides* (Tinoco et al., 2010) and the barley powdery mildew fungus *Blumeria graminis* (Nowara et al., 2010). Subsequent reports demonstrated the functionality of HIGS in suppressing diseases caused by the fungal pathogens *Puccinia* spp. (Yin et al., 2011; 2015; Zhang et al., 2012; Panwar et al., 2013), *Fusarium* spp. (Koch et al., 2013; Ghag et al., 2014; Cheng et al., 2015; Hu et al., 2015; Chen et al., 2016), *Sclerotinia sclerotiorum* (Andrade et al., 2015) and *Rhizoctonia solani* (Zhou et al., 2016), as well as by the oomycete pathogens *Phytophthora infestans* (Jahan et al., 2015; Sanju et al., 2015) and *Bremia lactucae* (Govindarajulu et al., 2015). Many of these pathogens make very intimate contact with host cells, potentially facilitating the occurrence of HIGS. In this study, we assessed whether HIGS can be used to suppress Verticillium wilt disease in tomato and *A. thaliana* by targeting previously identified virulence factors of *V. dahliae*, a pathogen that is known not to invade host cells at any stage of disease development.

## RESULTS

### *Tobacco rattle virus*-based silencing in tomato compromises *V. dahliae Ave1* expression

Tobacco rattle virus (TRV)-based virus-induced gene silencing (VIGS) has extensively been used in various plant species, including tomato (Liu et al., 2002; Senthil-Kumar et al., 2007), and TRV-based VIGS has successfully been used to investigate candidate genes for their involvement in Verticillium wilt resistance in tomato (Fradin et al., 2009). In order to investigate whether HIGS can be established against the xylem-colonizing fungus *V. dahliae* that is not known to penetrate adjacent living cells, we attempted to exploit TRV-based VIGS to produce dsRNAs that are targeted towards *V. dahliae Ave1* transcripts. The experiment was performed in *Ve1* tomato plants that are normally immune to infection by *Ave1*-carrying *V. dahliae* strains, such that successful HIGS would immediately result in vascular wilt disease that does not occur if *Ave1* expression is not compromised (Fradin et al., 2009; de Jonge et al., 2012). To this end, a 1:1 mixture of *Agrobacterium tumefaciens* cultures carrying *TRV1* and *TRV2::Ave1* (Figure 1A) was infiltrated into cotyledons of *Ve1* tomato plants. A recombinant construct containing a fragment of the *GUS* gene (*TRV2::GUS*) was used as a negative control (Figure 1A). At ten days after TRV treatment, plants were challenged with either the *V. dahliae* race 1 strain JR2 (Faino et al., 2015), or an *Ave1* deletion mutant (*V. dahliae* JR2*△Ave1*; de Jonge *et al*., 2012), and inspected for Verticillium wilt symptoms (stunting and wilting) up to 14 days post inoculation (dpi). As expected, no significant disease symptoms were observed on *TRV::GUS*-treated plants that were inoculated with the wild-type race 1 *V. dahliae* strain (Figure 2A), indicating that TRV treatment by itself does not compromise Ave1-triggered immunity in *Ve1* tomato plants. Furthermore, the *Ave1* deletion mutant caused clear Verticillium wilt disease, as Verticillium wilt disease developed on *Ve1* plants treated with *TRV2::Ave1* and subsequent inoculation with the *Ave1* deletion strain (Figure 2A). However, intriguingly, Verticillium wilt disease also developed on *Ve1* plants upon *TRV2::Ave1* treatment and subsequent inoculation with the wild-type race 1 *V. dahliae* strain (Figure 2A). This finding suggests that *Ave1* expression in *V. dahliae* is indeed compromised due to TRV-induced HIGS in tomato. The compromised immunity was confirmed by fungal recovery assays by plating stem sections on potato dextrose agar (PDA) plates, and by fungal biomass quantification in stem sections of the inoculated plants (Figure 2).

**Figure 1.**
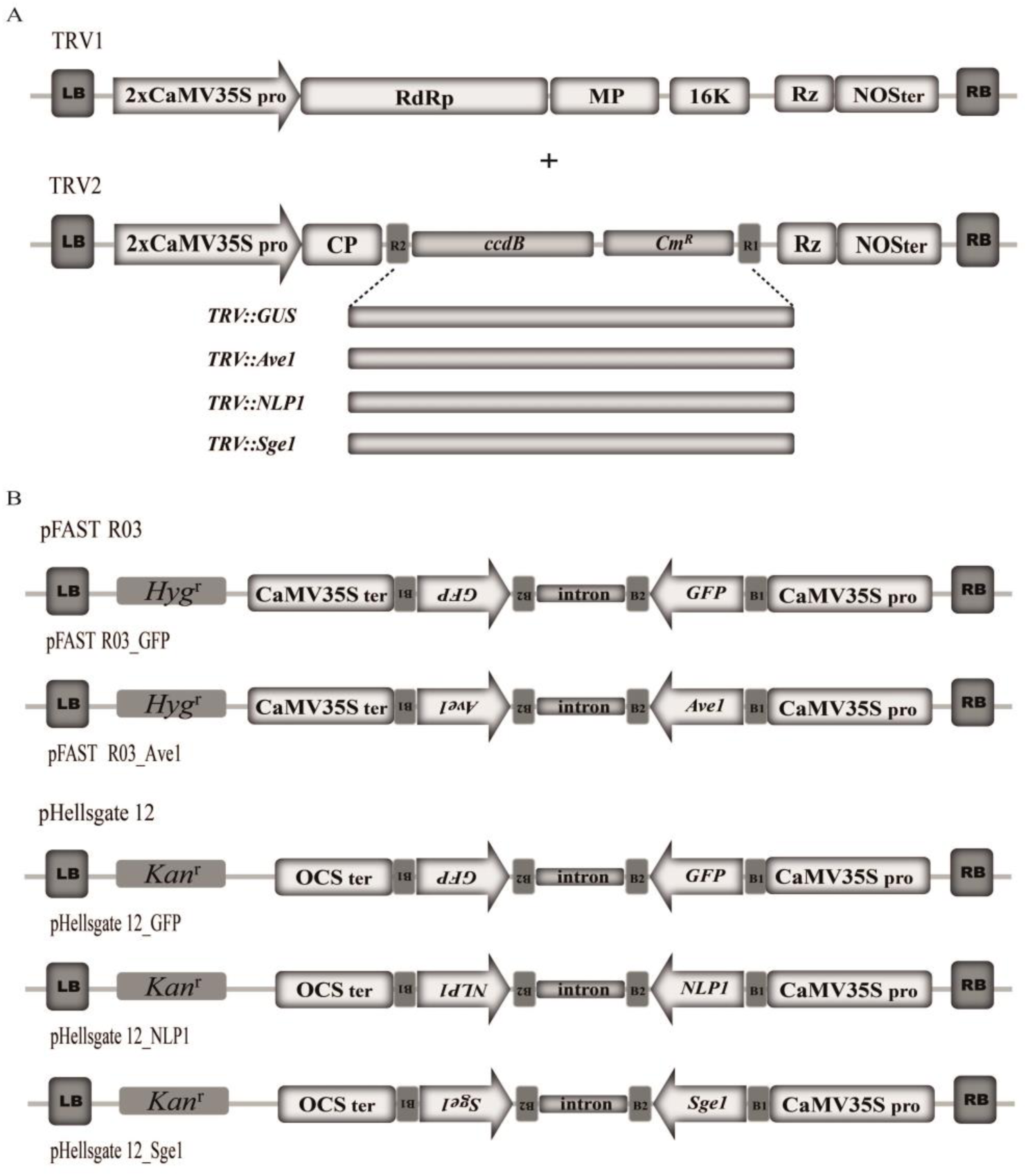
Schematic organization of the T-DNA region of the binary vectors used for gene silencing. (A) Schematic representation of the T-DNA region of the *Tobacco rattle virus* (TRV)-based virus-induced gene silencing (VIGS) vectors. *Verticillium dahliae Ave1*, *Sge1* and *NLP1* DNA fragments were inserted between the double CaMV35S promoter (2X35 CaMV35Spro) and the nopaline synthase gene terminator (NOSter) in the TRV2 vector to generate the VIGS vectors *TRV::Ave1*, *TRV::Sge1* and *TRV::NLP1*, respectively. Control construct *TRV::GUS* was described earlier (Song et al., 2016). RdRp, RNA-dependent RNA polymerase; 16K, 16 kDa cysteine-rich protein; MP, movement protein; CP, coat protein; Rz, self-cleaving ribozyme; *ccdB*, negative selection marker used in bacteria; *Cm*^*R*^, chloramphenicol resistance marker; R1 and R2, *att*R1 and *att*R2 sites. (B) Schematic diagrams of the T-DNA region of the binary vectors generated for producing a hairpin RNA of *Verticillium* genes *Ave1* (pFAST R03_Ave1), *NLP1* (pHellsgate 12_NLP1) and *Sge1* (pHellsgate 12_Sge1), as well as the *green fluorescent protein* gene (pFAST R03_GFP and pHellsgate 12_GFP) in transgenic *A. thaliana* plants. CaMV35Spro, CaMV35S promoter; CaMV35Ster, CaMV35S terminator; OCSter, octopine synthase gene terminator; *Hyg*^*r*^, hygromycin resistance gene; *Kan*^*r*^, kanamycin resistance gene; B1 and B2, *att*B1 and *att*B2 sites. LB and RB, left borders of T-DNA.

**Figure 2.**
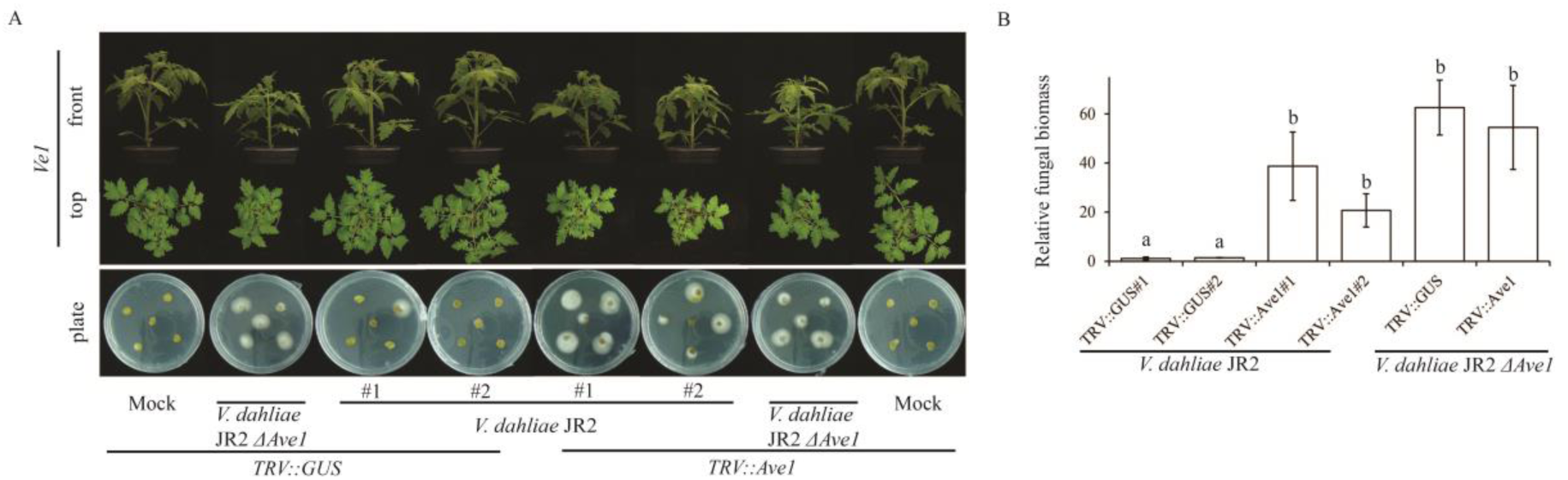
TRV-mediated fungal gene silencing in tomato plants compromises *Verticillium dahliae Ave1* expression. (A) Upon inoculation with the wild-type race 1 *V. dahliae* strain JR2, the impairment of Ave1-triggerred immunity in *Ve1* tomato plants treated with *TRV::Ave1* when compared with the *TRV::GUS*-treated plants evidenced by stunted *Ve1* plants at 14 days post inoculation (14 dpi) and fungal outgrowth upon plating stem sections on potato dextrose agar (PDA). The *Ave1* deletion mutant (*V. dahliae* JR2 *ΔAve1*) was used as *Verticillium* inoculation control. Plants were photographed at 14 dpi. (B) Fungal biomass determined with real-time PCR in *Verticillium*-inoculated *Ve1* plants at 14 dpi. Bars represent *Verticillium ITS* levels relative to tomato *actin* levels (for equilibration) with standard deviation in a sample of three pooled plants. The fungal biomass in *Ve1* tomato plants upon *TRV::GUS* treatment and subsequent inoculation with the wild-type race 1 *V. dahliae* strain is set to 1. Different letter labels indicate significant differences (*P* < 0.05).

It was recently demonstrated that *Tobacco mosaic virus* (TMV) may infect fungi in addition to plants, remaining for up to six subcultures, and also persisted in plants infected by the virus-infected fungus (Mascia et al., 2014). This finding raises the theoretical possibility that also TRV may infect *V. dahliae* and cause VIGS (directly) rather than HIGS from inside the tomato cells (indirectly). To exclude that the impairment of Ave1-triggered immunity in *Ve1* tomato plants is due to TRV-infection of *V. dahliae* itself, stem sections from *Verticillium*-inoculated *TRV::GUS*- and *TRV::Ave1*-treated tomato plants were placed on PDA plates. A single colony that grew from wild-type *V. dahliae*-inoculated *TRV::GUS*-treated plants (*V. dahliae* JR2^*TRV::GUS*^) and *Ave1* deletion mutant-inoculated *TRV::Ave1*-treated plants (*V. dahliae* JR2*△Ave1^TRV::Ave1^*), and three independent colonies that grew from wild-type *V. dahliae*-inoculated *TRV::Ave1*-treated plants (*V. dahliae* JR2^*TRV::Ave1*^) were subjected to PCR to detect a viral coat protein gene fragment of TRV (*TRV2_CP*), showing that *TRV2_CP* was not detected in all fungal isolates (Figure S1A). Furthermore, the fungal isolates were used to infect *Ve1* tomato plants and tomato plants that lack *Ve1*. This analysis showed that, similar to *V. dahliae* JR2^*TRV::GUS*^, also *V. dahliae* JR2^*TRV::Ave1*^ induced no disease symptoms on tomato plants expressing *Ve1*, while tomato plants lacking *Ve1* showed clear Verticillium wilt disease (Figure S1B-D). These data support the hypothesis that the impairment of Ave1-triggered immunity in *Ve1* plants is not caused by TRV-infection of *V. dahliae*, but genuinely by HIGS through TRV-treatment of tomato.

### TRV-based fungal gene silencing in tomato inhibits Verticillium wilt disease

To further investigate the potential of TRV-mediated HIGS against *V. dahliae* in tomato, two previously identified virulence genes of *V. dahliae* were targeted. The first target gene is *NLP1*, encoding a member of the necrosis- and ethylene-inducing-like protein (NLP) family in *V. dahliae*, and targeted deletion of *NLP1* in *V. dahliae* significantly compromises virulence on tomato as well as *A. thaliana* plants (Santhanam et al., 2013). The second candidate gene is *V. dahliae Sge1*, encoding a homolog of the transcription factor Sge1 (SIX Gene Expression 1) in *F. oxsporum*, and *V. dahliae* mutants of the *Sge1* are non-pathogenic on tomato (Santhanam and Thomma, 2013). To produce dsRNAs of the gene fragments *in planta*, cotyledons of ten-day-old Moneymaker tomato plants were treated with the silencing constructs *TRV::GUS*, *TRV2::NLP1* and *TRV2::Sge1* in combination with *TRV1* (Figure 1A), respectively. At ten days after TRV treatment, plants were challenged with either the *V. dahliae* strain JR2 (Faino et al., 2015), a *NLP1* deletion mutant (*V. dahliae* JR2*△NLP1*; Santhanam et al., 2013), or a *Sge1* deletion mutant (*V. dahliae* JR2*△Sge1*; Santhanam and Thomma, 2013), and monitored for Verticillium wilt symptoms on tomato plants at 14 dpi. As expected, significantly compromised Verticillium wilt symptoms were observed on Moneymaker tomato plants upon *TRV2::GUS* or *TRV2::NLP1* treatment and subsequent inoculation with the *NLP1* deletion mutant of *V. dahliae* strain JR2 (Figure 3A). Interestingly, upon inoculation with the wild-type *V. dahliae* strain JR2, a moderate reduction of Verticillium wilt symptoms was observed on Moneymaker tomato plants treated with *TRV2::NLP1* when compared to *TRV::GUS*-treated plants (Figure 3A). The plants that were treated with *TRV::NLP1* and subsequent inoculation with the wild-type *V. dahliae* strain JR2 showed reduced Verticillium wilt symptoms but were not as diseased as plants upon inoculation with the *NLP1* deletion mutant or water (Figure 3A). These data are further supported by fungal biomass quantifications in stem sections of the inoculated plants (Figure 3B). In contrast, no significant Verticillium wilt disease reduction was observed in Moneymaker tomato plants upon the *TRV::Sge1* treatment and subsequent inoculation with the wild-type *V. dahliae* strain JR2, although fungal biomass quantifications revealed that less fungal biomass accumulated *in planta* in the *TRV::Sge1*-treated plants than the *TRV::GUS*-treated plants followed by inoculation with the wild-type *V. dahliae* strain JR2 (Figure 4A; B). To determine whether TRV-mediated targeting transcripts of *V. dahliae Sge1* in tomato results in complete silencing of the *V. dahliae Sge1* gene, we performed real-time PCR to measure relative expression level for the *Sge1* gene in *V. dahliae* JR2 inoculating with the *TRV::Sge1*-treated plants compared to the *TRV::Sge1*-treated plants. However, only a slight reduction in *Sge1* expression in *TRV::Sge1*-targeted *V. dahliae* was monitored when compared with *TRV::GUS*-targeted *V. dahliae* (Figure 4C). In conclusion, although not all TRV-based RNAi constructs targeting *V. dahliae* transcripts in tomato suppressed Verticillium wilt disease, TRV-mediated transient HIGS against *V. dahliae* in tomato can be achieved.

**Figure 3.**
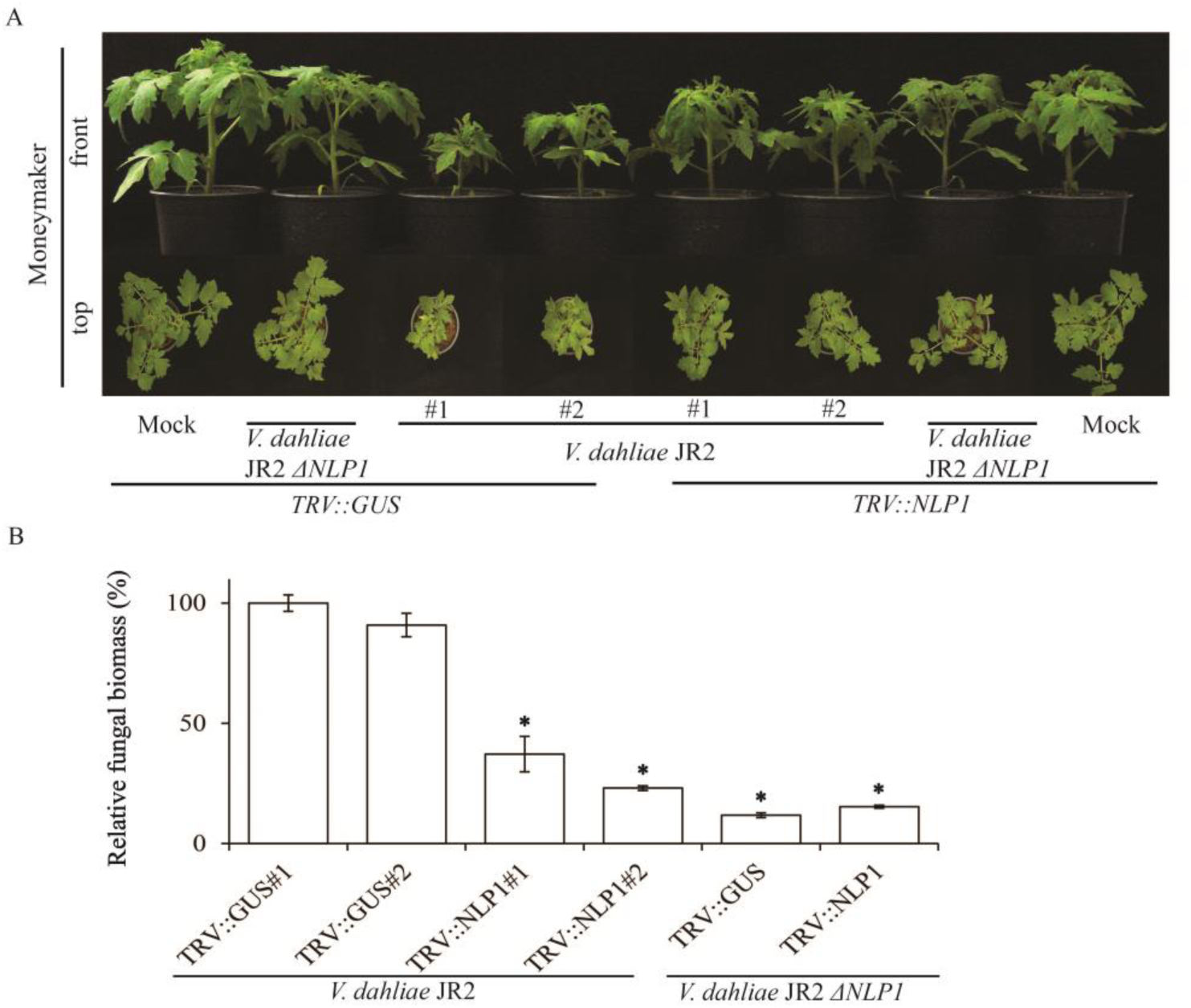
Effect of TRV-mediated *NLP1* silencing in Moneymaker tomato plants on *Verticillium dahliae* inoculation. (A) Agroinfiltration with the *TRV::NLP1* construct resulted in the suppression of Verticillium wilt symptoms on tomato plants, whereas no effect on disease development was observed on plants treated with the *TRV::GUS*. The *NLP1* deletion mutant (*V. dahliae* JR2 *ΔNLP1*) was used as *Verticillium* inoculation control. Plants were photographed at 14 dpi. (B) Fungal biomass determined with real-time PCR in *Verticillium*-inoculated Moneymaker tomato plants at 14 dpi. Bars represent *Verticillium ITS* levels relative to tomato *actin* levels (for equilibration) with standard deviation in a sample of three pooled plants. The fungal biomass in tomato plants upon *TRV::GUS* treatment and subsequent inoculation with the wild-type *V. dahliae* strain JR2 is set to 100% (control). Asterisks indicate significant differences when compared with the *TRV::GUS*-treated plants upon inoculation with the *V. dahliae* strain JR2 (*P* < 0.05).

**Figure 4.**
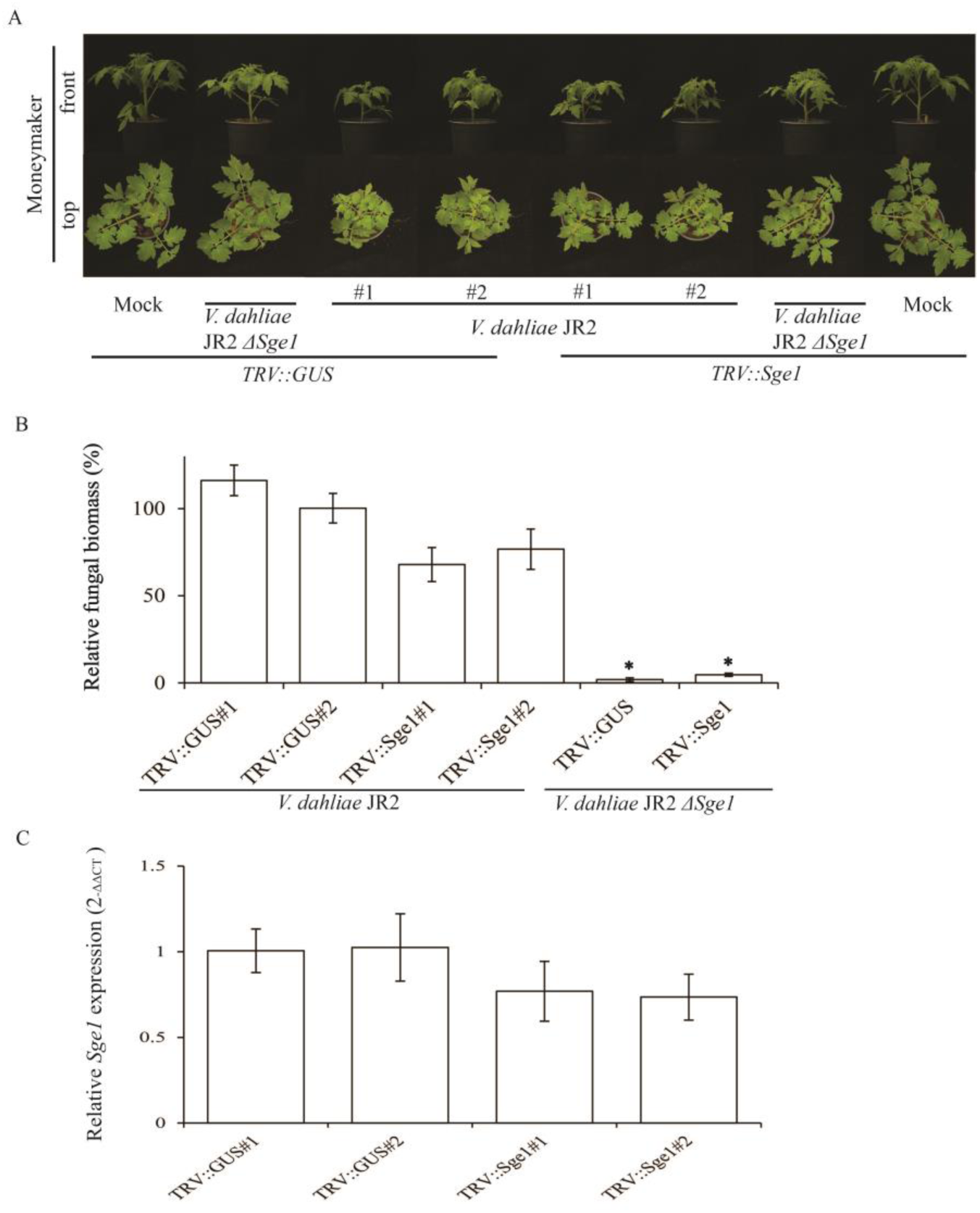
Effect of TRV-mediated *Sge1* silencing in tomato plants on *Verticillium dahliae* inoculation. (A) Upon inoculation with the *V. dahliae* strain JR2, no effect on disease development was observed on *TRV::Sge1*-treated plants compared to *TRV::GUS-*treated plants. The *Sge1* deletion mutant (*V. dahliae* JR2 *ΔSge1*) was used as *Verticillium* inoculation control. Plants were photographed at 14 dpi. (B) Fungal biomass determined with real-time PCR in *Verticillium*-inoculated Moneymaker tomato plants at 14 dpi. Bars represent *Verticillium ITS* levels relative to tomato *actin* levels (for equilibration) with standard deviation in a sample of three pooled plants. The fungal biomass in tomato plants upon *TRV::GUS* treatment and subsequent inoculation with the wild-type *V. dahliae* strain JR2 is set to 100% (control). Asterisks indicate significant differences when compared with the *TRV::GUS*-treated plants upon inoculation with the *V. dahliae* strain JR2 (*P* < 0.05). (C) Relative expression level for the *Sge1* gene was determined by using qRT-PCR at 14 days post inoculation with the wild-type *V. dahliae* strain on *TRV::Sge1*-, and *TRV::GUS*-treated plants. Bars represent levels of *Sge1* transcripts relative to the transcript levels of *V. dahliae GAPDH* (GAPDH, glyceraldehyde-3-phosphate dehydrogenase; for normalization) with standard deviation of a sample of three pooled plants. *Sge1* expression in *V. dahliae* the *TRV::GUS*-treated plants upon inoculation the wild-type strain *V. dahliae* is set to 1.

### HIGS in *Ve1*-transgenic *A. thaliana* does not impair Ave1-triggerred immunity

To assess whether HIGS against *V. dahliae* can be made operational in stable transgenic plants by expressing dsRNAs, we exploited hairpin RNA-based RNAi to produce dsRNAs to target *V. dahliae Ave1* transcripts in *Ve1*-expressing *A. thaliana* plants. To this end, a fragment of the *V. dahliae Ave1* gene was cloned into the Gateway vector pFAST R03 (Shimada et al., 2010) to obtain the RNAi construct pFAST R03_Ave1 (Figure 1B) that leads to hairpin RNA formation after transcription. A recombinant RNAi construct containing a fragment of the *green fluorescent protein* (*GFP*) gene (pFAST R03_GFP) was used as a negative control (Figure 1B). Subsequently, the *Ave1* and *GFP* RNAi constructs were transformed into recipient *Ve1*-expressing *A. thaliana* plants (Fradin et al., 2011) (Figure S2A). No obvious developmental alterations were observed in the transgenic plants when compared with the recipient Col-0 and *Ve1* plants (Figure 5A), and three independent *Ave1* RNAi lines expressing *Ve1* (pFAST R03_Ave1 in *Ve1*-1, pFAST R03_Ave1 in *Ve1*-2 and pFAST R03_Ave1 in *Ve1*-3) as well as transgenic and non-transgenic control lines were inoculated with either the *V. dahliae* race 1 strain JR2 or an *Ave1* deletion mutant, and monitored for Verticillium wilt symptoms up to 21 dpi. As expected, upon mock-inoculation or inoculation with the *V. dahliae* JR2, no disease symptoms were observed in *Ve1* plants and *GFP*-RNAi *Ve1* plants (Figure 5A). In contrast, *GFP*- or *Ave1*-RNAi Col-0 plants lacking *Ve1* were as diseased as non-transformed control lines (Figure 5A). However, despite transcripts for hairpin *Ave1* formation in *Ave1*-RNAi *Ve1* plants were detected (Figure S2A), Verticillium wilt symptoms were not observed in *Ave1*-RNAi *Ve1* plants upon inoculation with the wild-type race 1 *V. dahliae* strain JR2 in repeated assays, while the *Ave1* deletion strain caused clear Verticillium wilt symptoms on *Ve1* plants (Figure 5A). The phenotypes correlated with the degree of *V. dahliae* colonization as determined with real-time PCR (Figure 5). These data show that expression of an RNAi construct targeting *Ave1* transcripts in *A. thaliana* plants expressing *Ve1* does not compromise Ave1-triggered immunity.

**Figure 5.**
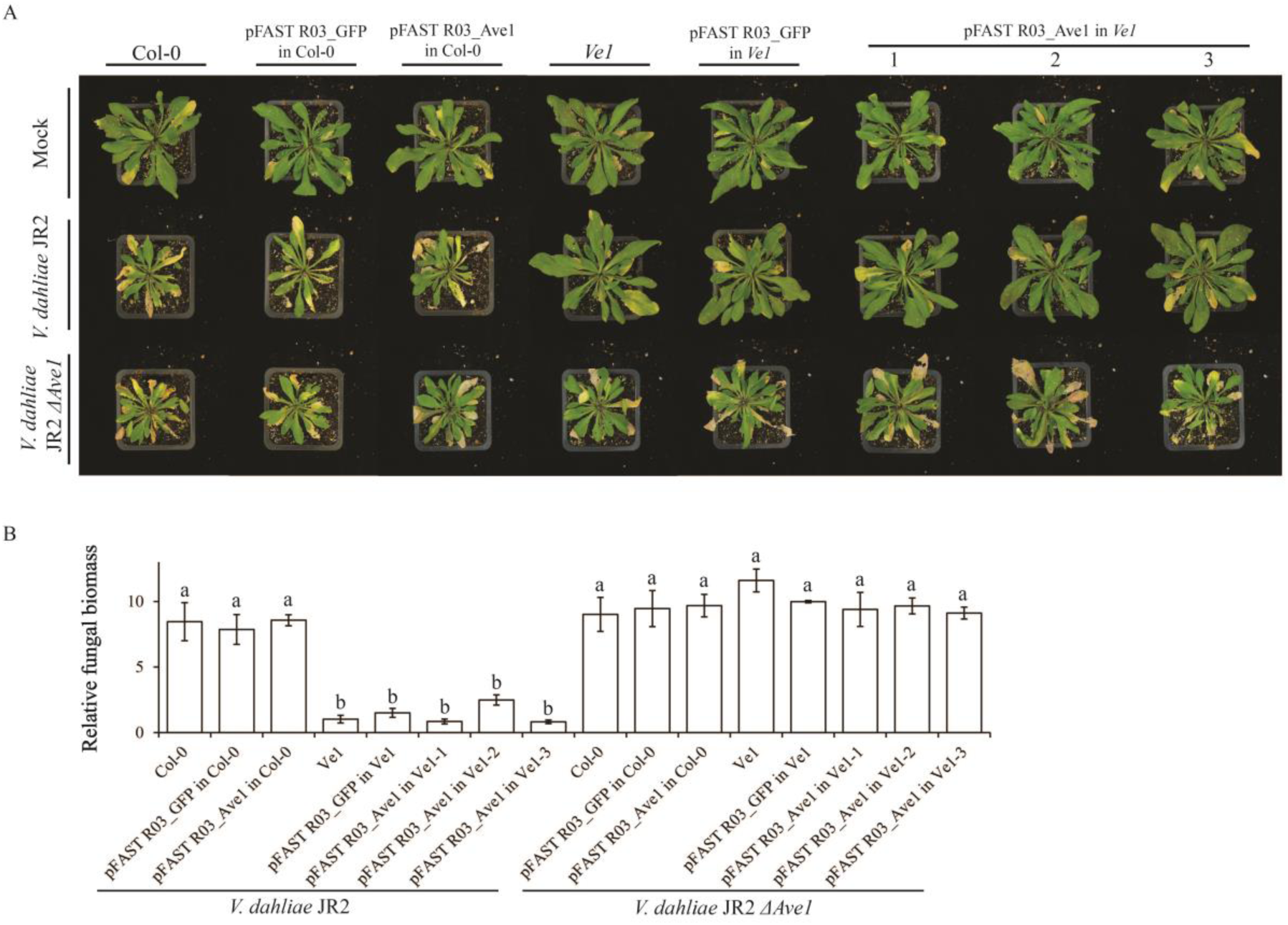
Analysis of *Ve1 Arabidopsis thaliana* plants expressing RNAi *Ave1* construct. (A) Upon inoculation with the race 1 *V. dahliae* strain JR2, *Ve1* plants expressing RNAi *Ave1* or *GFP* construct do not show Verticillium wilt symptoms, whereas typical Verticillium wilt symptoms are recorded on plants with or without tomato *Ve1* upon inoculation with either the *V. dahliae* JR2 or *V. dahliae* JR2 *ΔAve1* at 21 dpi. Col-0 plants with or without tomato *Ve1* were used as a controls.The *V. dahliae* JR2 *ΔAve1* strain was used as *Verticillium* inoculation control. (B) Fungal biomass determined with real-time PCR in *Verticillium*-inoculated *Arabidopsis* plants at 21 dpi. Bars represent *Verticillium ITS* levels relative to *AtRuBisCo* (RuBisCo, ribulose-1, 5-bisphoshate-carboxylase/oxygenase) levels (for equilibration) with standard deviation in a sample of five pooled plants. The fungal biomass in *Ve1* plants upon inoculation with the wild-type race 1 *V. dahliae* strain JR2 is set to 1 (control). Three independent lines carrying the pFAST R03_Ave1 construct are shown (1, 2 and 3). Different letter labels indicate significant differences (*P* < 0.05).

### HIGS in *A. thaliana* can reduce Verticillium wilt

To further investigate whether HIGS against *V. dahliae* can be established in stable transgenic *A. thaliana* plants by expressing dsRNAs, *V. dahliae NLP1* and *Sge1* were targeted. To this end, RNAi constructs pHellsgate 12_NLP1 and pHellsgate 12_Sge1 were generated (Figure 1B). The recombinant RNAi construct carrying a fragment of the *GFP* gene (pHellsgate 12_GFP) was used as a negative control (Figure 1B). Subsequently, RNAi constructs targeting *NLP1*, *Sge1*, or *GFP* were transformed into *A. thaliana* ecotype Col-0, and independent *NLP1*-, *Sge1*, or *GFP*-RNAi lines were selected (Figure S2B; C). No phenotypic alterations were observed in *NLP1*- or *Sge1*-RNAi plants when compared with the recipient *A. thaliana* Col-0 plants or *GFP*-RNAi plants (Figure 6A; 7A). Three independent *NLP1*-RNAi lines (pHellsgate 12_*NLP1*-1, pHellsgate 12_*NLP1*-2 and pHellsgate 12_*NLP1-*3) as well as *GFP*-RNAi and non-transgenic control lines were assayed for the development of Verticillium wilt symptoms. As expected, markedly compromised Verticillium wilt symptoms were observed on *A. thaliana* plants upon inoculation with the *NLP1* deletion mutant (Figure 6A). Interestingly, upon inoculation with the *V. dahliae* JR2, a significant reduction of Verticillium wilt symptoms was observed in *NLP1*-RNAi plants when compared with *GFP*-RNAi and non-transgenic controls (Figure 6A). These data are further supported by fungal biomass quantifications in stem sections of the inoculated plants (Figure 6B). Additionally, three independent *Sge1*-RNAi lines (pHellsgate 12_*Sge1*-1, pHellsgate 12_*Sge1*-2 and pHellsgate 12_*Sge1-*3) as well as *GFP*-RNAi and non-transformed control lines were assayed for Verticillium wilt disease development. Intriguingly, we observed a marked reduction of Verticillium wilt symptoms in *Sge1*-RNAi *A. thaliana* lines inoculated with *V. dahliae* JR2 (Figure 7A). In contrast, *GFP*-RNAi *A. thaliana* lines were as susceptible as non-transgenic control lines (Figure 7A). The phenotypes correlated with the level of *V. dahliae* biomass as determined with real-time PCR (Figure 7). Collectively, these results suggest that, although not all RNAi constructs targeting *V. dahliae* transcripts in *A. thaliana* induced HIGS against *V. dahliae*, hairpin RNA-mediated HIGS in *A. thaliana* can reduce Verticillium wilt disease.

**Figure 6.**
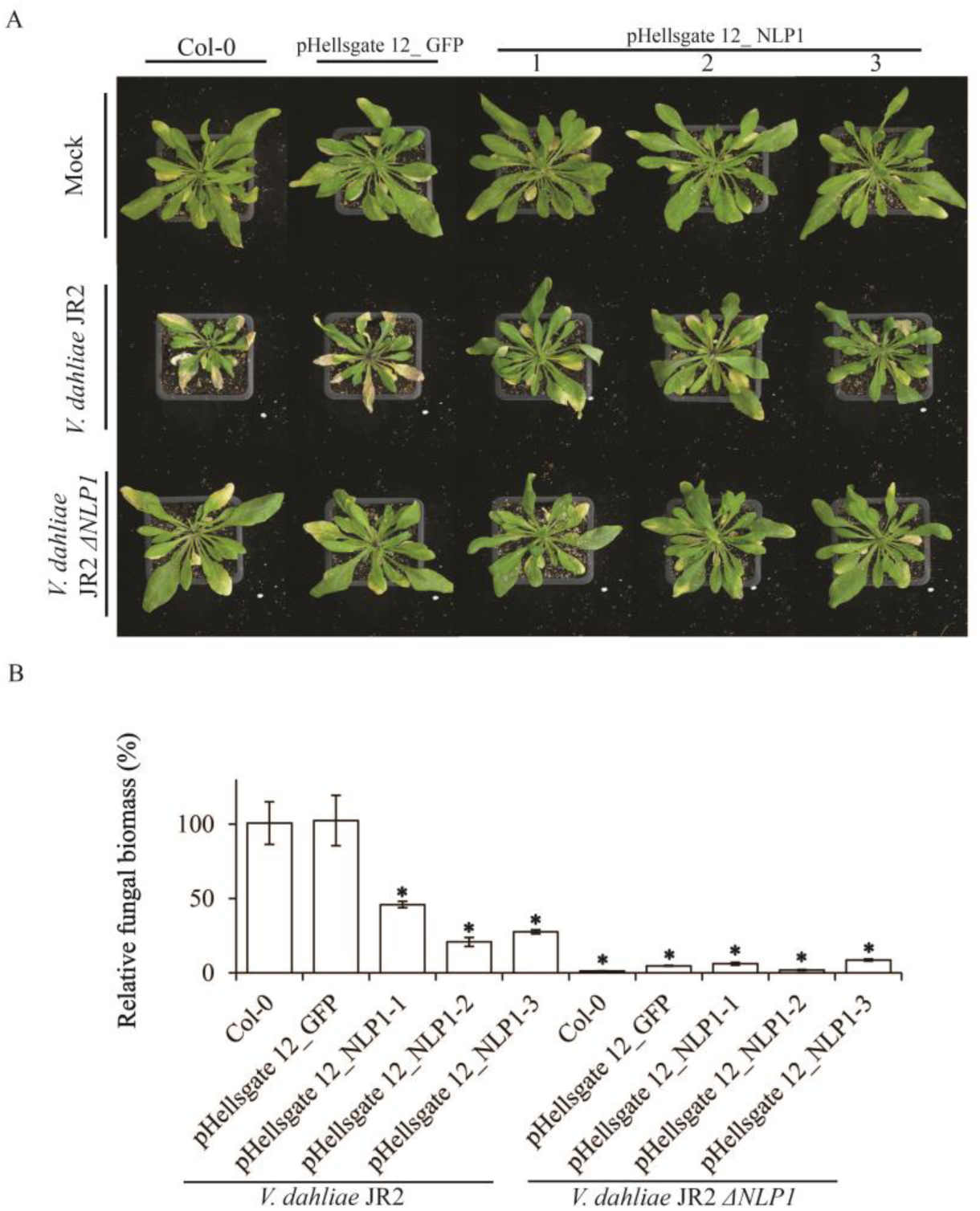
*Arabidopsis thaliana* Col-0 plants expressing *NLP1* RNAi construct provide resistance against *Verticillium dahliae*. (A) Typical appearance of non-transgenic *A. thaliana* and transgenic lines carrying the pHellsgate 12_NLP1 construct to target *NLP1* transcripts upon mock-inoculation or inoculation with *V. dahliae* strain JR2 or *V. dahliae* JR2 *ΔNLP1* at 21 dpi. (B) Fungal biomass determined with real-time PCR in *Verticillium*-inoculated *Arabidopsis* plants at 21 dpi. Bars represent *Verticillium ITS* levels relative to *AtRuBisCo* (RuBisCo, ribulose-1, 5-bisphoshate-carboxylase/oxygenase) levels (for equilibration) with standard deviation in a sample of five pooled plants. The fungal biomass in Col-0 is set to 100% (control). Three independent lines carrying the pHellsgate 12_NLP1 construct are shown (1, 2 and 3). Asterisks indicate significant differences when compared with Col-0 (*P* < 0.05).

**Figure 7.**
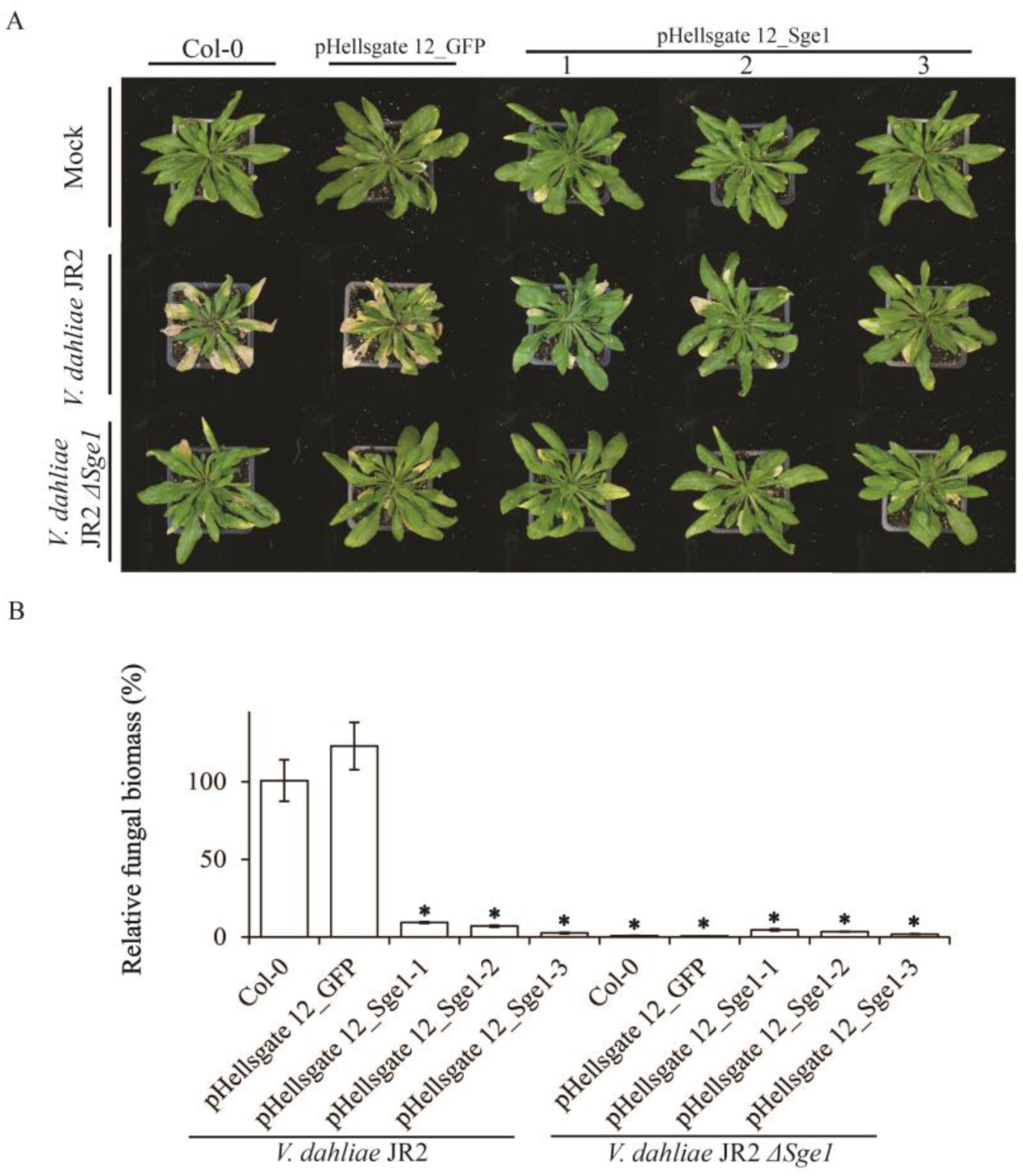
*Arabidopsis thaliana* Col-0 plants expressing *Sge1* RNAi construct confer resistance against *Verticillium dahliae*. (A) Typical appearance of non-transgenic *A. thaliana* and transgenic lines harboring the pHellsgate 12_Sge1 construct to target *Sge1* transcripts upon mock-inoculation or inoculation with *V. dahliae* strain JR2 or *V. dahliae* JR2 *ΔSge1* at 21 dpi. (B) Fungal biomass determined with real-time PCR in *Verticillium*-inoculated *Arabidopsis* plants at 21 dpi. Bars represent *Verticillium ITS* levels relative to *AtRuBisCo* (RuBisCo, ribulose-1, 5-bisphoshate-carboxylase/oxygenase) levels (for equilibration) with standard deviation in a sample of five pooled plants. The fungal biomass in Col-0 is set to 100% (control). Three independent lines carrying the pHellsgate 12_Sge1 construct are shown (1, 2 and 3). Asterisks indicate significant differences when compared with Col-0 (*P* < 0.05).

## DISCUSSION

In this manuscript, we show that HIGS against *V. dahliae* can be achieved through TRV-based fungal gene silencing in tomato, and through hairpin RNA-mediated fungal gene silencing in stable transgenic *A. thaliana* lines. We established the TRV-mediated HIGS assay through targeting *V. dahliae Ave1* transcripts in *Ve1* tomato plants, and further used this approach to assess whether HIGS against *V. dahliae* in tomato can be established through TRV constructs targeting previously identified *V. dahliae* virulence factors. We also investigated whether HIGS against *V. dahliae* can be established in transgenic *A. thaliana* plants through hairpin RNA-based RNAi constructs targeting transcripts of the same previously identified *V. dahliae* virulence genes. Our results clearly show that plants transiently (in tomato) or stably (in *A. thaliana*) expressing RNAi constructs targeting transcripts of genes that are essential for *V. dahliae* pathogenicity can become protected from Verticillium wilt disease. Our results are in line with, and extend beyond, recent reports on protection of cotton plants stably expressing an RNAi construct against *V. dahliae* (Zhang et al., 2016), and on bidirectional cross-kingdom RNAi and fungal uptake of external RNAs to confer plant protection (Wang et al., 2016).

Reports on HIGS against fungal pathogen infections have accumulated over recent years (Yin et al., 2011; 2015; Zhang et al., 2012; Koch et al., 2013; Panwar et al., 2013; Ghag et al., 2014; Andrade et al., 2015; Cheng et al., 2015; Hu et al., 2015; Chen et al., 2016; Zhou et al., 2016). Among these reports, disease suppression was observed upon silencing of various types of genes, including those that encode the biosynthesis of structural components such chitin and ergosterol, but also genes involved in developmental regulation, secondary metabolism and pathogenicity. Therefore, the selection of suitable target genes is arguably the most important prerequisite for developing a successful HIGS against fungal pathogens. We selected HIGS target genes based on our previous studies of gene deletion mutants in *V. dahliae* with significantly compromised virulence (*NLP1*, Santhanam et al., 2013; and *Sge1*, Santhanam and Thomma, 2013). On beforehand, it was not clear whether silencing of such genes rather than gene deletion would lead to visible virulence phenotypes, as the protein encoded by the target gene may not be completely absent. Indeed, TRV-mediated HIGS of *Sge1* in tomato did not lead to compromised Verticillium wilt symptoms, which may be explained by the incomplete silencing of the *Sge1* gene in *V. dahliae* (Figure 4C). This also explains why RNAi-mediated HIGS of *V. dahliae Ave1* did not compromise *Ve1*-mediated immunity in transgenic *A. thaliana* plants. Also in the cases where the expected visual phenotypes were obtained, fungal biomass quantifications revealed that more fungal biomass accumulated in the inoculated plants upon HIGS of the fungal target gene than upon inoculation with the corresponding *V. dahliae* deletion mutant (Figure 3B; Figure 4B; Figure 6B; Figure 7B). Thus, arguably, RNAi may not be appropriate to target genes of which the activity is required early in the infection process when RNAi may not have taken its full effect, or genes of which a low-dose of transcripts is biologically active.

Mobility of small RNAs within organisms is a well-known phenomenon, facilitating gene silencing in adjacent cells and surrounding or even distant tissues (Weiberg et al., 2015). Over recent years, several examples of exchange of small RNAs between host plants and invading pathogens have been described, although the mechanistic details of the actual exchange remains to be elucidated (Knip et al., 2014). Nevertheless, small RNA-based bidirectional cross-kingdom gene silencing has been proposed as a common mechanism for cross-kingdom gene regulation in plant-pathogen interactions (Weiberg et al., 2015; Chaloner et al., 2016; Wang et al., 2016). For example, endogenous small RNAs from the fungus *Botrytis cinerea* have been proposed to transfer into host plants to target defense-related plant transcripts to promote disease development (Weiberg et al., 2013). In this manner, HIGS taps into a process that naturally occurs between plants and pathogens. A search for pathogen-derived small RNAs matching transcripts of host plants or plant-derived small RNAs targeting transcripts of the invading pathogens may facilitate the development of HIGS strategies to engineer resistance in plants against pathogens for which no natural resistance sources have been identified.

## EXPERIMENTAL PROCEDURES

### Plant growth conditions and manipulations

Plants were grown at 21°C/19°C during 16 h/8 h light/dark photoperiods, respectively, in the climate chamber or the greenhouse with a relative humidity of ~75%, and 100 W⋅m^-2^ supplemental light when light intensity dropped below 150 W⋅m^-2^. *A. thaliana* transformations were performed as described (Clough and Bent, 1998).

#### Generation of the constructs

The Gateway-compatible *Tobacco rattle virus* (TRV) two-component *Agrobacterium*-mediated expression system was used for gene silencing in tomato as previously described (Liu et al., 2002), while the Gateway-compatible vectors pFAST R03 (Shimada et al., 2010) and pHellsgate 12 (Helliwell and Waterhouse, 2003) for hairpin RNA-mediated gene silencing were used to generate stable Arabidopsis transformants. The three *V. dahliae* genes *Ave1* (de Jonge et al., 2012), *NLP1* (Santhanam et al., 2013) and *Sge1* (Santhanam and Thomma, 2013) were selected for RNAi-based HIGS. Gene annotations for *V. dahliae Ave1*, *Sge1* and *NLP1* were obtained from Ensembl Genomes database (http://fungi.ensembl.org/Verticillium_dahliaejr2/Info/Index). Selected DNA fragments were amplified by PCR from the corresponding plasmids using gene-specific primers listed in Table S1. The DNA fragments were cloned into pDONR207 by using the Gateway^®^ BP Clonase^®^ II Enzyme Mix (Invitrogen, California, USA) to generate entry vectors, and all the entry vectors were verified by sequencing. Subsequently, the entry vector pDONR207 carrying the *Ave1* fragment was transferred to TRV2 and pFAST R03 to generate constructs *TRV2::Ave1* and pFAST R03_Ave1 (Figure 1), while pDONR207 entry vectors carrying *NLP1* or *Sge1* fragment were recombined into pTRV2 and pHellsgate 12 to generate constructs *TRV2::NLP1*, *TRV2::Sge1*, pHellsgate 12_NLP1 and pHellsgate 12_Sge1 (Figure 1) by using Gateway^®^ LR Clonase^®^ II Enzyme Mix (Invitrogen, California, USA). All constructs were transformed to *Agrobacterium tumefaciens* strain GV3101 (pMP90) by electroporation.

#### TRV treatment

TRV vectors were agroinfiltrated as previously described (Liu et al., 2002; Fradin et al., 2011). Briefly, cotyledons of 10-day-old tomato (*Solanum lycopersicum* cv. Moneymaker (*ve1*) or *35S::Ve1* tomato (*Ve1*); Fradin et al., 2009) were infiltrated as 1:1 mixtures of *pTRV1* and *pTRV2* constructs. 10-15 days after TRV inoculation, plants were inoculated with race 1 *V. dahliae* strain JR2 (Faino *et al*., 2015); the corresponding mutants: *V. dahliae* JR2 *ΔAve1* (de Jonge et al., 2012); *V. dahliae* JR2 *ΔNLP1* (Santhanam et al., 2013); *V. dahliae* JR2 *ΔSge1* (Santhanam and Thomma, 2013); or tap water as control. The inoculated plants were evaluated by observing disease symptoms up to 14 days post inoculation (dpi).

#### Verticillium wilt disease and fungal recovery assays

*V. dahliae* was grown on potato dextrose agar (PDA) at 22 °C, and conidia were collected from 7- to 10-day-old plates and washed with tap water. Disease assays on tomato plants were performed as previously described (Fradin et al., 2009). Briefly, twenty-day-old *Ve1* tomato plants (for *Verticillium* inoculation after TRV inoculation) or ten-day-old *Ve1* and *ve1* tomato plants (for inoculation with *Verticillium* colonies re-isolated from infected tomato plants) were uprooted, the roots were rinsed in water, dipped for 5 min in a suspension of 10^6^ conidiospores/mL water, and transplanted to soil. *Verticillium* outgrowth assays of *Ve1* tomato plants, canopy area measurement and fungal biomass quantification in tomato plants were performed as previously described (Fradin et al., 2009; Santhanam et al., 2013). *Verticillium* disease assay on *A. thaliana*, as well as fungal biomass quantification in infected *A. thaliana* plants were performed as previously described (Ellendorff et al., 2009; Song et al., 2016). The fungus-specific primer ITS1-F, based on the internal transcribed spacer (ITS) region of the ribosomal DNA, in combination with the *V. dahliae*-specific reverse primer ST-Ve1-R (Ellendorff et al., 2009) were used to measure fungal colonization. Primers for tomato *actin* and *A. thaliana RuBisCo* (Table S1) were used as endogenous plant control.

#### Quantitative Real Time-PCR (qRT-PCR) and Reverse transcription-PCR (RT-PCR)

To determine expression of *V. dahliae Sge1* gene for silencing, stems of TRV-treated tomato plants were harvested at 14 days post *Verticillium* inoculation as described above, and flash frozen in liquid nitrogen, and stored at -80 °C for total RNA isolation.

To check *Ave1*, *NLP1*, *Sge1*, *GFP* DNA fragment presence in the transcripts of the corresponding transgenic *A. thaliana* lines, two-week-old transgenic and non-transgenic *A. thaliana* lines were harvested and ground into a powder in liquid nitrogen. Total RNA extraction, cDNA synthesis and RT-PCR were performed as described earlier (Song et al., 2016). Primers for hairpin expression analysis are listed in Table S1. To analyze expression of *Sge1* gene for silencing, qRT-PCR was conducted by using primers Sge1-F(qRT) and Sge1-R(qRT) with *V. dahliae GAPDH* as an endogenous control (Table S1) as previously described (Santhanam and Thomma, 2013).

## ACKNOWLEDGEMENTS

Y.S. acknowledges a PhD fellowship from the China Scholarship Council (CSC). B.P.H.J.T. is supported by a Vici grant of the Research Council for Earth and Life sciences (ALW) of the Netherlands Organization for Scientific Research (NWO). Constantinos Patinios and Yidong Wang are acknowledged for technical assistance and Bert Essenstam for excellent plant care at Unifarm. The authors declare no conflict of interest.

---

**Figure S1.**
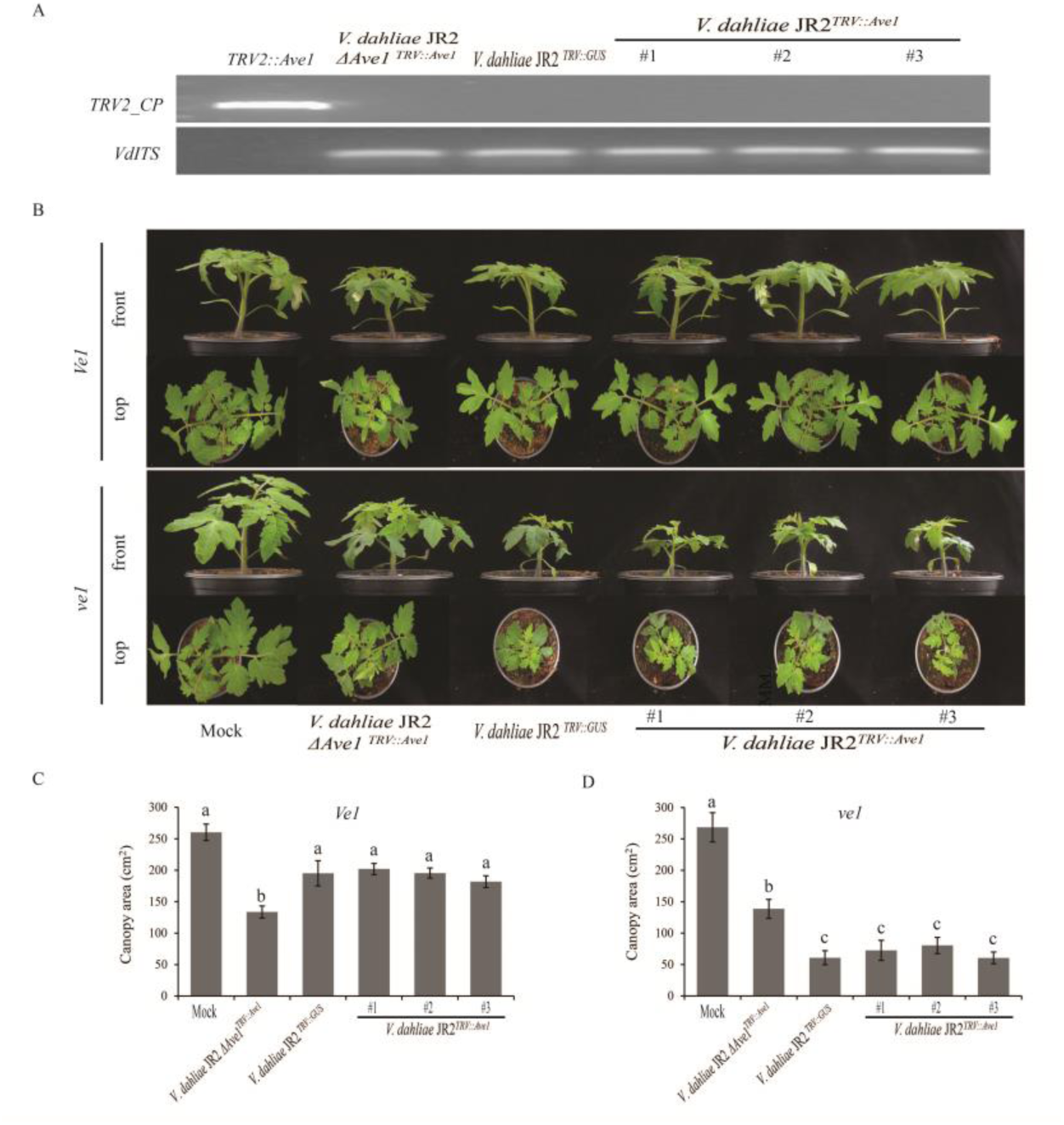
Analysis of *Verticillium dahliae* colonies re-isolated from TRV-treated *Ve1* tomato plants. (A) Viral coat protein gene fragment (*TRV2_CP*) was not detected by PCR in re-isolated *V. dahliae* colonies *V. dahliae* JR2 *∆Ave1^TRV::Ave1^*, *V. dahliae* JR2 ^*TRV::GUS*^ and *V. dahliae* JR2^*TRV::Ave1*^ (#1, #2 and #3) that grew from wild-type *V. dahliae*-inoculated *TRV::GUS*-treated plants, *Ave1* deletion strain-inoculated *TRV::Ave1*-treated plants and wild-type *V. dahliae*-inoculated *TRV::Ave1*-treated plants, respectively. Construct *TRV::Ave1* was used as a positive control. As an endogenous control, a fragment of the *V. dahliae ITS* was amplified from all re-isolated *V. dahliae* colonies. (B) Typical appearance of *Ve1* tomato plants (*Ve1*) and Moneymaker tomato plants lacking *Ve1* (*ve1*) upon mock-inoculation or inoculation with re-isolated strains *V. dahliae* JR2 *∆Ave1^TRV::Ave1^*, *V. dahliae* JR2 ^*TRV::GUS*^, or *V. dahliae*JR2^*TRV::Ave1*^ at 14 dpi. (C) Average canopy area of ten plants inoculated with re-isolated strains described above or mock inoculation. Different letters indicate significant differences (*P* < 0.5).

**Figure S2.**
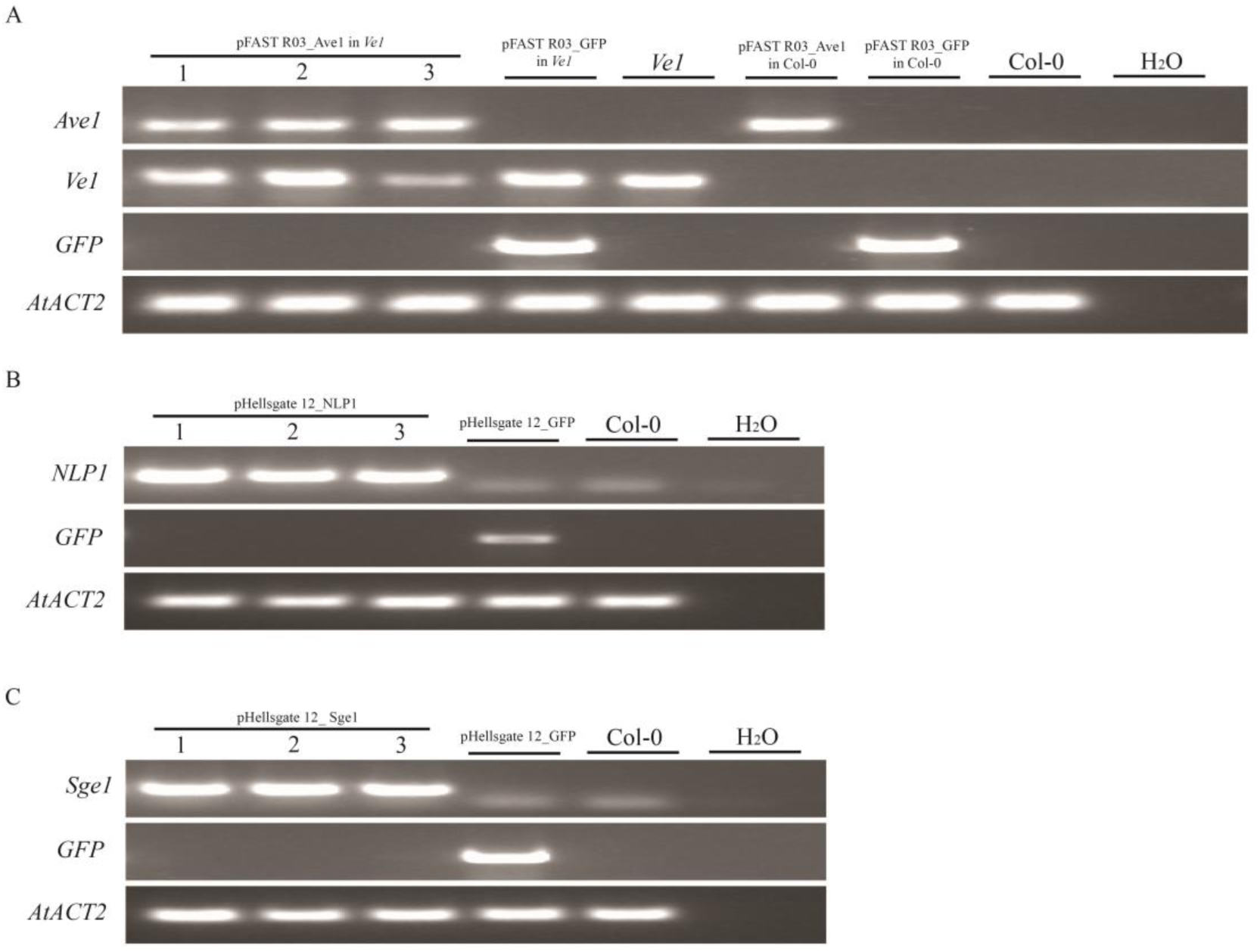
PCR amplification from cDNA of the corresponding transgenic *Arabidopsis thaliana* lines. (A) Transcripts for hairpin *Ave1* or *GFP* formation in transgenic lines were detected by reverse transcription-PCR (RT-PCR). For pFAST R03_Ave1 construct in *Ve1* plants three independent transgenic lines are shown (1, 2 and 3), while other lines expressing pFAST R03_Ave1 construct or pFAST R03_GFP construct are shown as controls. (B) Transcripts for hairpin *NLP1* or *GFP* formation in transgenic lines were PCR-detected. For pHellsgate 12_NLP1 construct in Col-0 plants three independent transgenic lines are shown (1, 2 and 3), and the corresponding control line expressing pHellsgate 12_GFP construct is shown. (C) Transcripts for hairpin *NLP1* or *GFP* formation in transgenic lines were PCR-detected. For pHellsgate 12_Sge1 construct in Col-0 plants three independent transgenic lines are shown (1, 2 and 3), and the corresponding control line expressing pHellsgate 12_GFP construct is shown. As an endogenous control, a fragment of the *AtACTIN2* gene was amplified from *A. thaliana* cDNA. *A. thaliana* Col-0 and water are used as PCR controls.

**Table S1.**
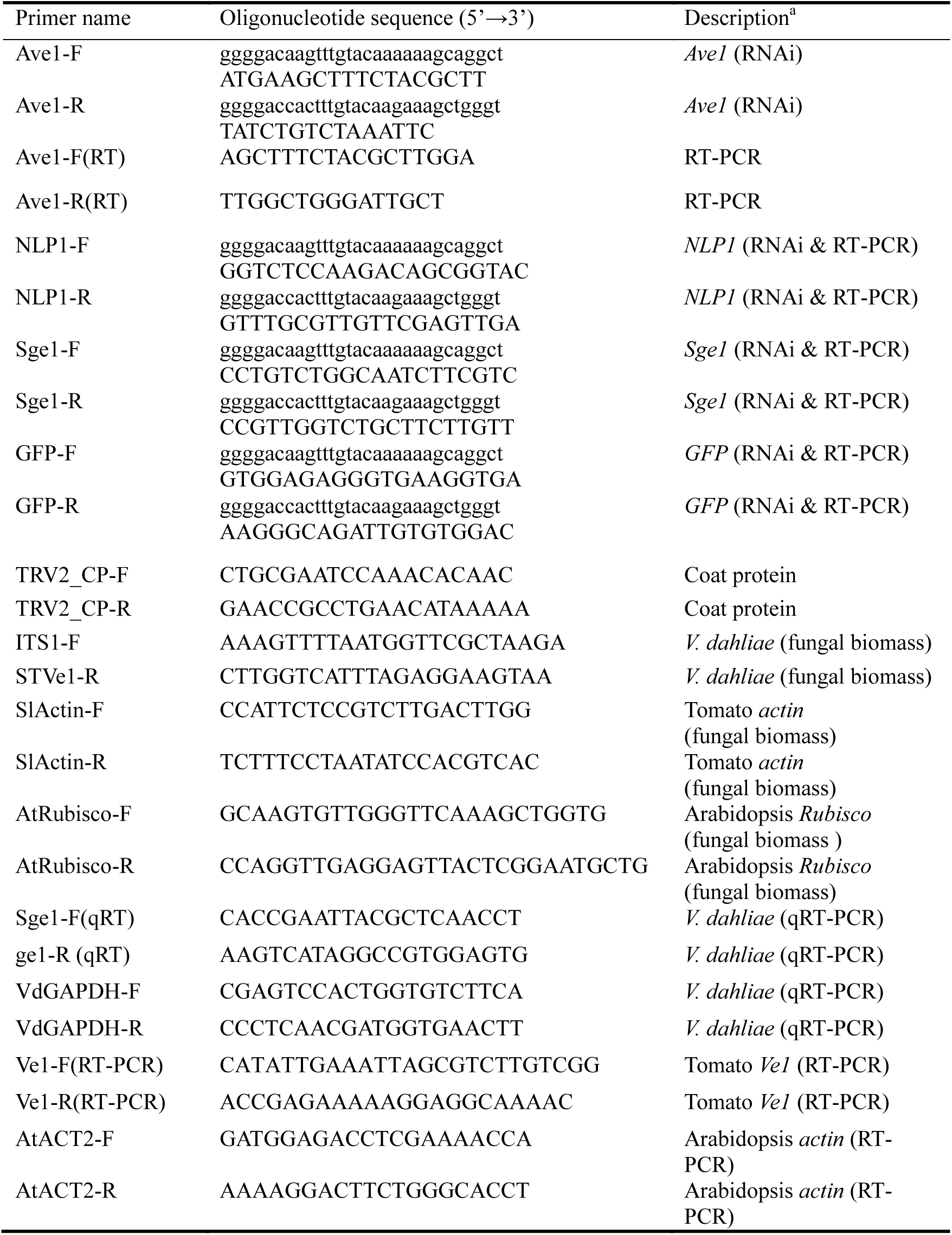
Primers used in this study.

